# Interdisciplinary and disciplinary identities: towards a theory of forms of knowledge change^1^

**DOI:** 10.1101/603449

**Authors:** Peter van den Besselaar

## Abstract

The concept of interdisciplinarity lacks theoretical understanding, and consequently, the number of indicators for interdisciplinarity is booming: out of field citations, betweenness centrality, or the set coherence, diversity, mediation, to mention a few. However, these indicators focus on characteristics of papers and journals, without referring clearly to the processes of knowledge production, communication and stabilization. Without understanding of the nature of interdisciplinarity and its various forms, the choice of indicators seems rather arbitrary. In this paper we argue that interdisciplinarity is one of the forms of development of research fields. We will show that interdisciplinarity is a temporary phase in knowledge dynamics. New fields emerge at the boundaries of existing fields as multidisciplinary research activities, and either develop through an interdisciplinary phase into a ‘new’ discipline’, or remain multidisciplinary for a while and generally disappear.

We use the analysis of journal-journal citation relations to map the identity and development of disciplinary, multidisciplinary and interdisciplinary research fields. A comparison results in three types of identities: (1) disciplinary fields, (2) interdisciplinarity as a developmental phase from multidisciplinary to disciplinary fields, and (3) unstable temporal multidisciplinary research topics at the boundaries between research fields. We conclude that the main difference between a mature interdisciplinary field and ‘normal’ disciplinary fields is that the former is newer than the latter. We propose to focus on change in the research landscape, instead of on indicators for what is only one (early or long term unstable) stage of interdisciplinarity. The mapping approach can be used to identify the nature of cognitive change in general, such as the emergence, growth, differentiation, merger, and decline of research fields.

## Introduction

Interdisciplinary always has been an issue for discussion, and a various definitions exist, all emphasizing different degrees of interdisciplinarity. The last decade, interest has been growing as science policy makers increasingly are inclined to stimulate interdisciplinary research (refs). The idea behind this is that innovation and fundamental break-throughs occurs not in the core of existing disciplines but between them: at the intersections were different disciplines meet.^2^

As a consequence of this, the request for indicators of interdisciplinarity comes up, as this would improve our ability to identify (new) interdisciplinary developments. The bibliometric community has responded to this, and we see a diversity of indicators being proposed. It started with out of field citations, and now is in the stage of more sophisticated network indicators. However, also the latter are generally based on the idea of out of field citations (e.g., Porter and Rafols 2009).

In this paper I aim to further develop another perspective on interdisciplinarity, using indicators for interdisciplinarity developed quite a while ago (Van den Besselaar, Heimeriks 2001). Basically I argue that we may better change the focus from interdisciplinarity to patterns of change – and interdisciplinarity only being one of these patterns. My argument consists of the following steps:

- Firstly, I’ll argue that interdisciplinarity is better conceptualized as a *position* in the scientific landscape, and not so much in terms of knowledge streams (*relations*) – although these are important too. Journal network structures do inform us about this.
- Secondly, if we look back to older interdisciplinary fields, we may learn more about the nature of interdisciplinarity than if we try to develop even more sophisticated indicators for something not very well understood. More specifically, this teaches that interdisciplinarity is a temporal stage in the development of new research fields, which either stabilize into a normal ‘disciplinary’ mode of communication, or remain unstable and ultimately disappear.
- Thirdly, if this is the case, we may better focus on *change* in the scientific landscape, and interdisciplinary integration is only one of the forms, next to specialization as the more general form of scientific development. But other forms of change also exist: birth of a new field, growth, decline, splitting (specialization), merging (integration), divergence, convergence and death.
- Fourthly, if these patterns of change do exist, and if the positional network structures are a correct representation, we may come up with theoretical understanding about the formal conditions under which these patterns of change occur. Sociology of science may inform us about the different social conditions for the emergence of the different patterns.

### Identifying interdisciplinarity

Following Price and many others, journals can be considered as the main carrier of scholarly communication, and this is still the case in the area of electronic access.^3^ In a previous paper (Van den Besselaar 2001), I argued that interdisciplinarity can straightforward be studied at the level of scholarly journals – as can be done for regular research fields (e.g., Doreain & Fararo 1985; Leydesdorff & Cozzens 1993; Van den Besselaar & Leydesdorff 1996).^4^ Research fields are represented as citation networks of journals. These networks have, as all communication networks, have two different structures: a positional structure and a relational structure.^5^

- Positional: journals with a similar citation behavior are clustered together: they have the same position in the network, without necessarily having strong citation relations (but often they do).
- Relational: journals with citation strong relations (mutually citing each other) are clustered together.

As already said, most indicators for interdisciplinarity take the relational network as frame of reference. The more a field cites outside its own journal set, the more interdisciplinary it is considered. However, theoretically, this is hardly convincing. If interdisciplinarity is mainly defined as theoretical and methodological integration, the use of knowledge from a different field is hardly an indicator. More explicitly, as we know, the more fundamental research fields are more cited than the application oriented ones. Both cite themselves, but the application oriented ones also heavily draw upon the fundamental ones for specific basic knowledge. That does not at all indicate that the plied fields are interdisciplinary.

In general, knowledge integration points at a different process. A field integrates knowledge from two or more fields, and therefore resembles to a certain extent these fields it integrates. Looking from the perspective of the fields that are integrated, the interdisciplinary field is to a certain extent similar, but not completely. This is captures by a positional analysis of journal-journal citation networks.

The analysis proceeds as follows.^6^ One or more core journals of the field under study are used as entrance. The citing environment of these journals is determined, and factor analyzed. Factors consist of journals with the same position in the network; that is journals with the same citing behavior. This indicates that they belong to the same research field. This is an old result which also can be used to study the dynamics of research fields over time (Van den Besselaar & Leydesdorff 1996) and for interdisciplinary fields (Van den Besselaar & Heimeriks 2001). Interdisciplinary journals are easily detected, as they load relatively high on more than one factor. This implies that they are similar to two or more research fields, an intuitively correct indicator for theoretical integration. Disciplinary research field, on the other hand, consist of journals that homogenously load only on one factor. A third category of journals are the multidisciplinary journals, that do not aim at integrating, but that cover more than one field – but have the same factor loadings pattern as the interdisciplinary journals. Examples are of course *Science, Nature, PNAS* covering all sciences, but these journals exist also at the level of disciplines, where they cover all or several fields (like *JACS* in chemistry, or *CACM* in computer science). In the next sections I will apply this approach on different interdisciplinary fields, of different ages, in order to investigate the nature of interdisciplinarity as one form of knowledge dynamics.

### Step 1: The communication structure of a traditional discipline – chemistry

Let’s start with a traditional discipline and show how the journal citation network of this discipline. We use the one the main^7^ journal in the field as entrance point: *Journal of the American Chemical Association*.^8^ We determine the environment^9^ of the *JACS*, which consists of 44 main chemistry journals. Factor-analyzing the journal-journal citation matrix in the citing dimension result in factors containing journals with similar citing behavior. These represent the different fields within chemistry. Figure 1 shows the results. The first factor is organic chemistry. Other factors are inorganic chemistry, chemical materials, physical chemistry, biochemistry, a physics factor focusing on materials, and finally macromolecules.

**Fig 1.**
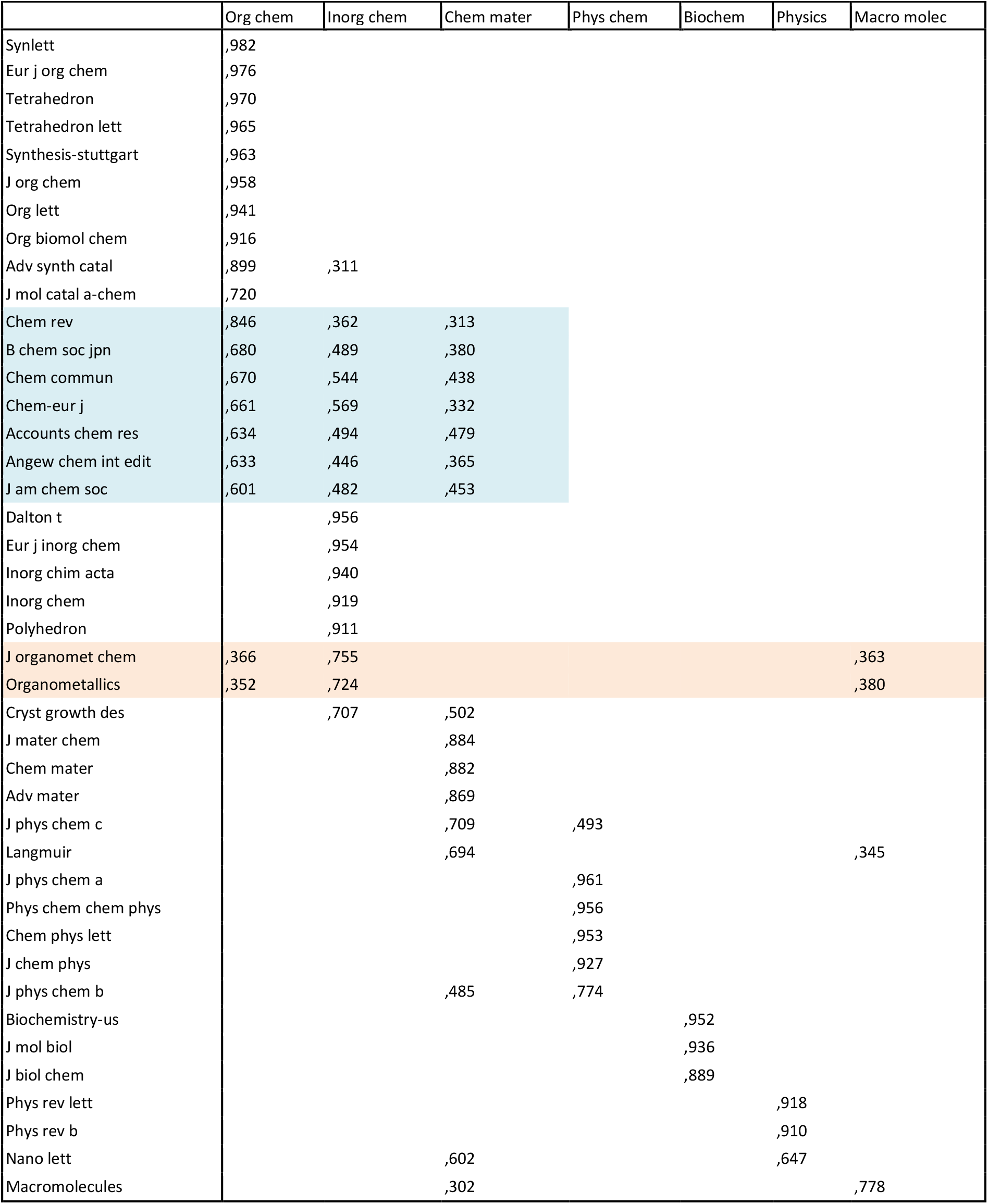
J Am Chem Soc (2007)

The main journals in the various chemical subfields load on one factor only, a typical disciplinary pattern. However, we find a few quite a few journals loading on more than one factor with a substantial (>0.3) loading: the hybrid borders of the various fields. However, within these journals we have to distinguish two categories. Firstly we find the journals within chemistry that cover various chemical subfields. These are generally high impact factor journals, and colored blue in the figure. These journals are covering more chemistry fields: the broad multidisciplinary journals, often the main journals of national chemical associations, or review journals. Secondly, we find journals with loadings on more factors: the interdisciplinary journals. These journals have a “double identity” as their citation pattern is similar to more than one research field, and obviously try to integrate different bodies of knowledge. In figure 1, this holds for example for the journals on organometals, operating at the boundaries of organic chemistry, inorganic chemistry and macromolecules research. Another example in figure 1 is *Nano Letters*, spanning the boundary between chemical materials and physical materials.

Many other fields can be used to exemplify the ‘normal’ case. Using a dynamic perspective, the factor structure over many years can be compared. Often they are rather stable over time, as well as the set of fields they relate to, as I showed elsewhere for a series of subdisciplines within chemistry.

Please note that we excluded *Science, Nature* and *PNAS* from the analyses presented here, as they make the picture less clear: these journals are multidisciplinary journals in the sense that they cover all important discoveries in all fields. This gives them a different role and function.

### Step 2: The communication structure of some old interdisciplinary fields

In the figure 1, an interesting observation can be made: an old interdisciplinary field, physical chemistry, actually shows a disciplinary factor structure. Most important journals load only on the physical chemistry factor itself, and only some interdisciplinary overlap exists – with chemical materials. Is this a normal pattern for established interdisciplinary fields? In order to answer the question, we now show the results for three of these old interdisciplinary fields: physical chemistry, biochemistry, and – to include also a social science example – social psychology.

Let’s start with physical chemistry. The main journal in the field is the *Journal of Chemical Physics*. This leads to a citation network of 35 journals, and using factor analysis we obtain the main components. As we now started with a core journal in physical chemistry, the number of physical chemistry journals in the set is larger than in the analysis of chemistry as a whole.

As already expected from figure 1, and confirmed by figure 2, also physical chemistry has a fairly disciplinary structure: the main journals in physical chemistry have a high loading on the factor representing the field, and only in the tail of the factors we find some journals loading also on a chemistry factor with mainly general chemistry journals – that also cover physical chemistry. In other words, physical chemistry may effectively have integrated physics and chemistry – the result is a new discipline with a citation structure as other disciplinary research fields.

**Figure 2:**
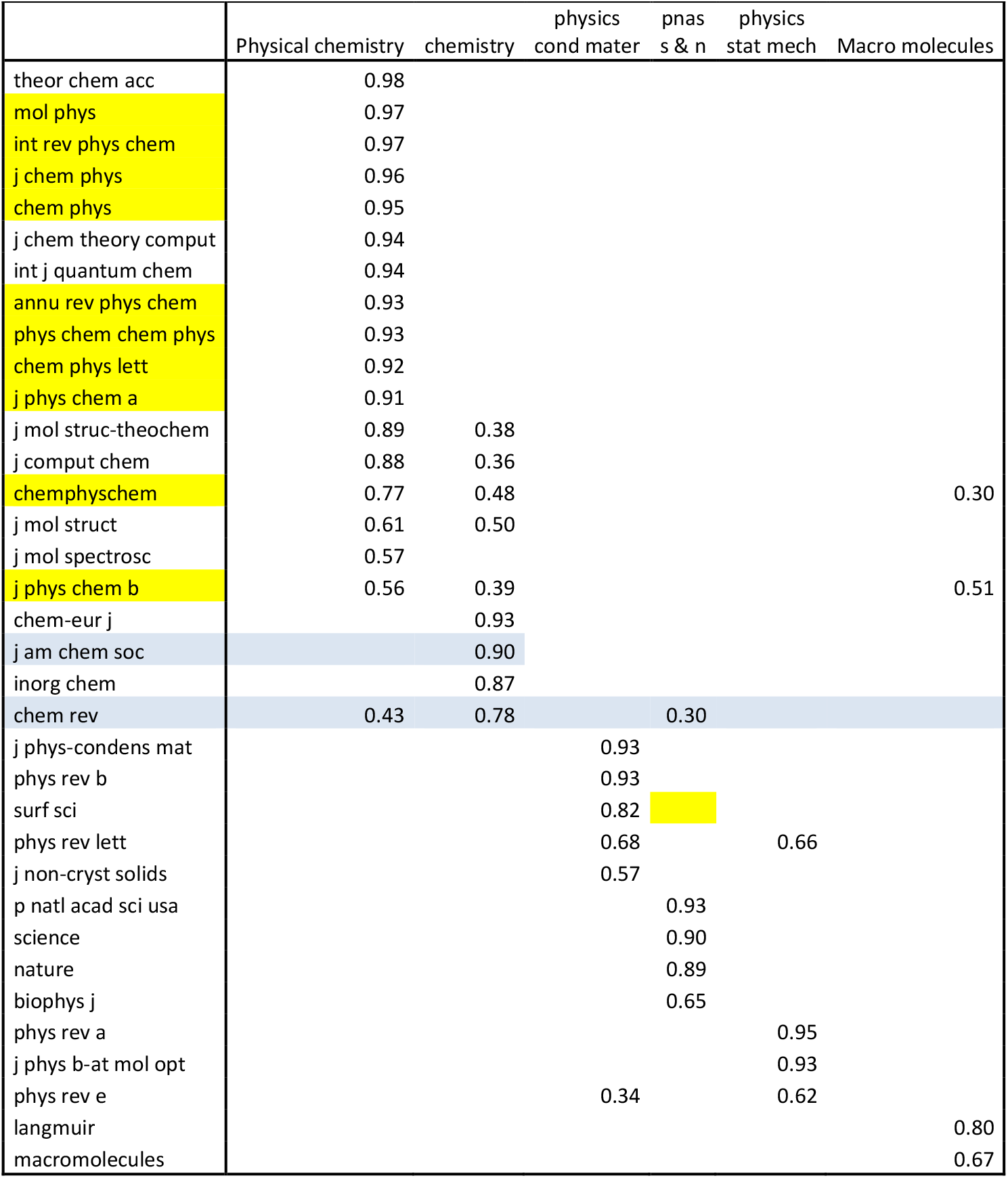
The factor structure of the Journal of *Chemical Physics*

The next example is biochemistry. The main journal is the *Journal of Biological Chemistry* with a 0.5% citation environment of 35 journals. Factor analyzing the matrix shows two interesting features. There is no other chemistry field visible than biochemistry. What once started at the border between chemistry and biology now is clearly part of biology. Second, the first three factors show overlap, suggesting a similarity between biochemistry, molecular biology and molecular cell biology. This opens the question to what extent this similarity is growing or not – that may give us an indicator for process of *convergence* and *divergence* within this set of three fields. This will be done in another paper.

**Fig 3:**
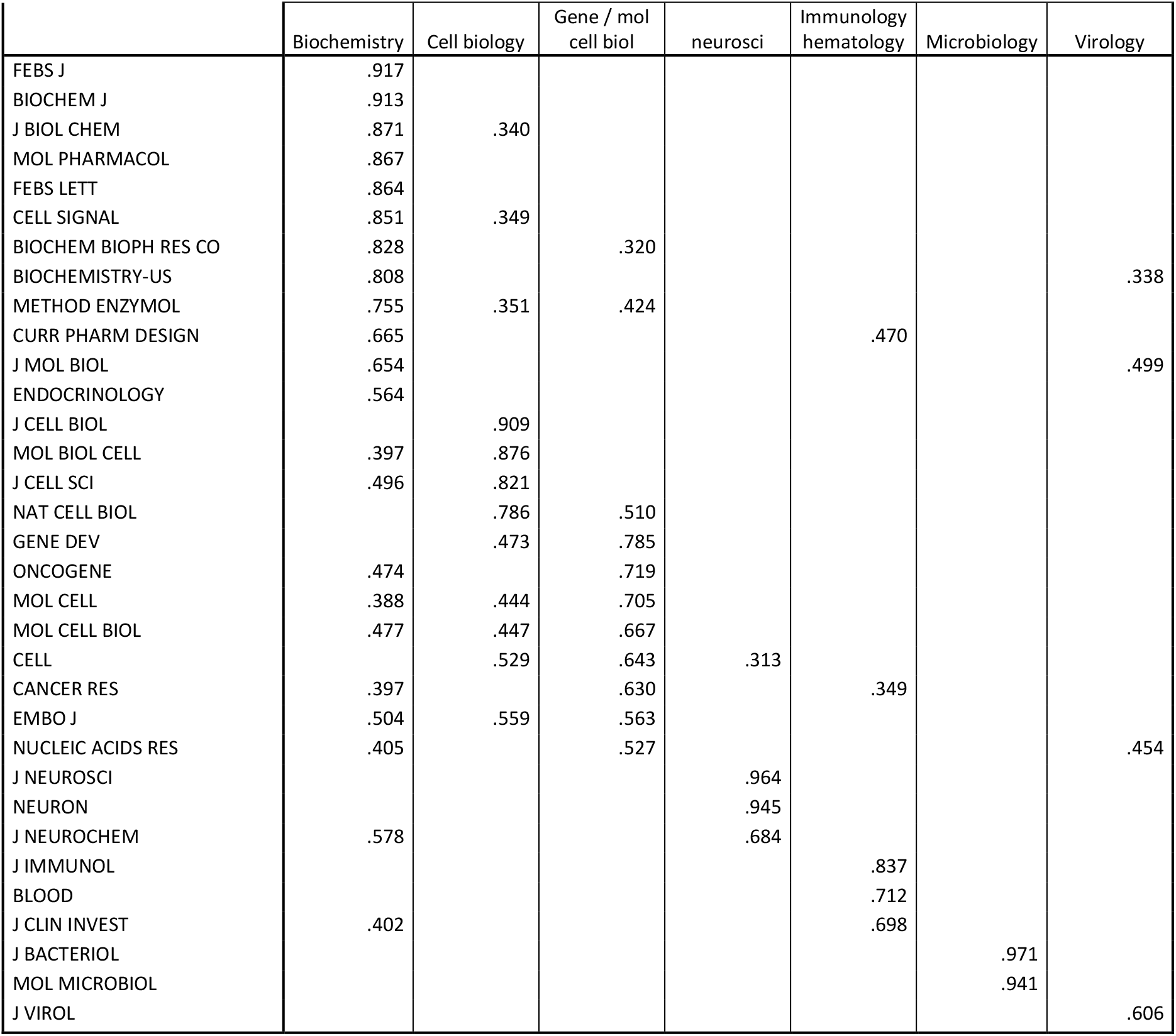
Journal of Biological Chemistry

The last example of an old interdisciplinary field discussed here is social psychology. We again use the most cited journal in the field as an entrance: the *Journal of Personal and Social Psychology*. This has a citation environment of 38 journals, of which many in social psychology. Even if the field may have emerged between sociology and psychology, there is nothing left of this anymore. The field has a clear disciplinary communication structure, with most of its journals only loading on the main social psychology factor. The other factors in the environment are all psychology. So emerging in the interdisciplinary space between sociology and psychology, the field has become disciplinary and converged with psychology (and diverged from sociology).

**Figure 4:**
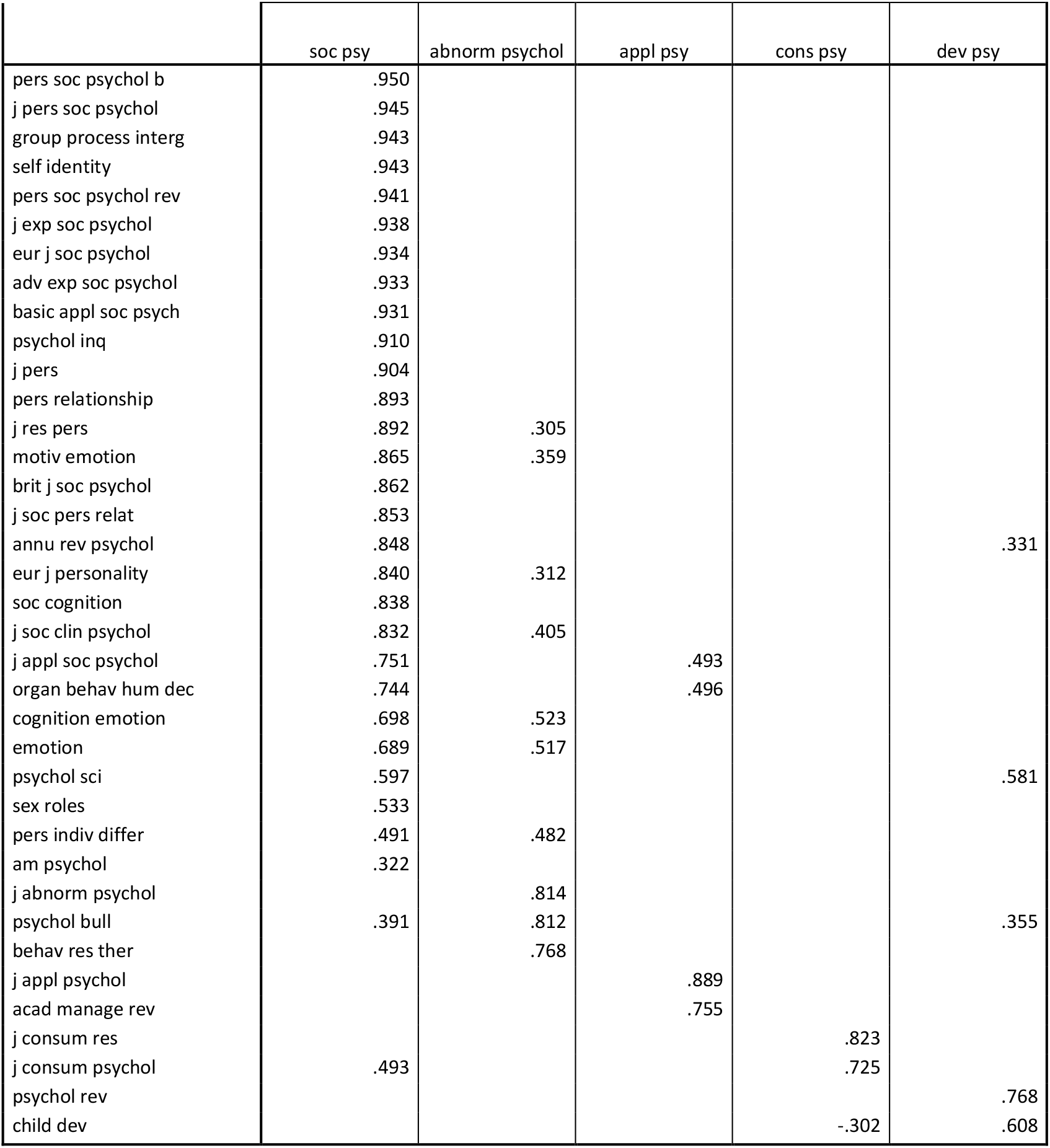
Citation environment of the *Journal of Personal and Social Psychology*

What do we learn from this analysis (Van den Besselaar, Heimeriks 2001)? Firstly, monodisciplinary fields are characterized by journals loading mainly on one factor that represents the field (e.g., the main fields within chemistry in table 1). Secondly, the journals that do load on more factors are often multidisciplinary journals covering all fields in a discipline (e.g., the general chemistry journals in table 1), or indicate the possible emergence of a multidisciplinary development (e.g., the organometallics journals in table 1). Finally, the ‘old examples’ of interdisciplinarity show that these fields in fact did become ‘disciplinary’: consisting of journals belonging one factor, that represents the field. In other words, they have a similar communication structure as the old disciplines. This is most strongly the case for social psychology, which has a very strong internal journal citation network, hardly linked to any other subfield – even within psychology. As we do not have the data to study their emergence, we now will have a closer look at some more recent examples of interdisciplinary fields, as this may teach us how this ‘disciplinary interdisciplinarity’ comes into existence.

## 3. The development of more recent interdisciplinary fields

Here we take up examples developed in previous publications (Van den Besselaar & Leydesdorff 1996; Van den Besselaar & Heimeriks 2001). In these papers we showed that the method introduced above can clearly be used to describe the development of research fields, and especially also for the development of interdisciplinary fields. Let’s have a look at two examples, artificial intelligence and cognitive science.

### 3.1 Artificial Intelligence: From interdisciplinarity to disciplinarity

An interesting type of development is seen in the case of Artificial Intelligence (AI). The field of AI started as a rather multidisciplinary field, as showed in table 5. The main journal in the field, *Artificial Intelligence Journal*, does not load on an AI factor, but is the third journal in a multidisciplinary factor with a cognitive science journal and a linguistics journal. The wider environment of *Artificial Intelligence Journal* consists of psychology, and a series of computer science related research fields.

**Figure 5:**
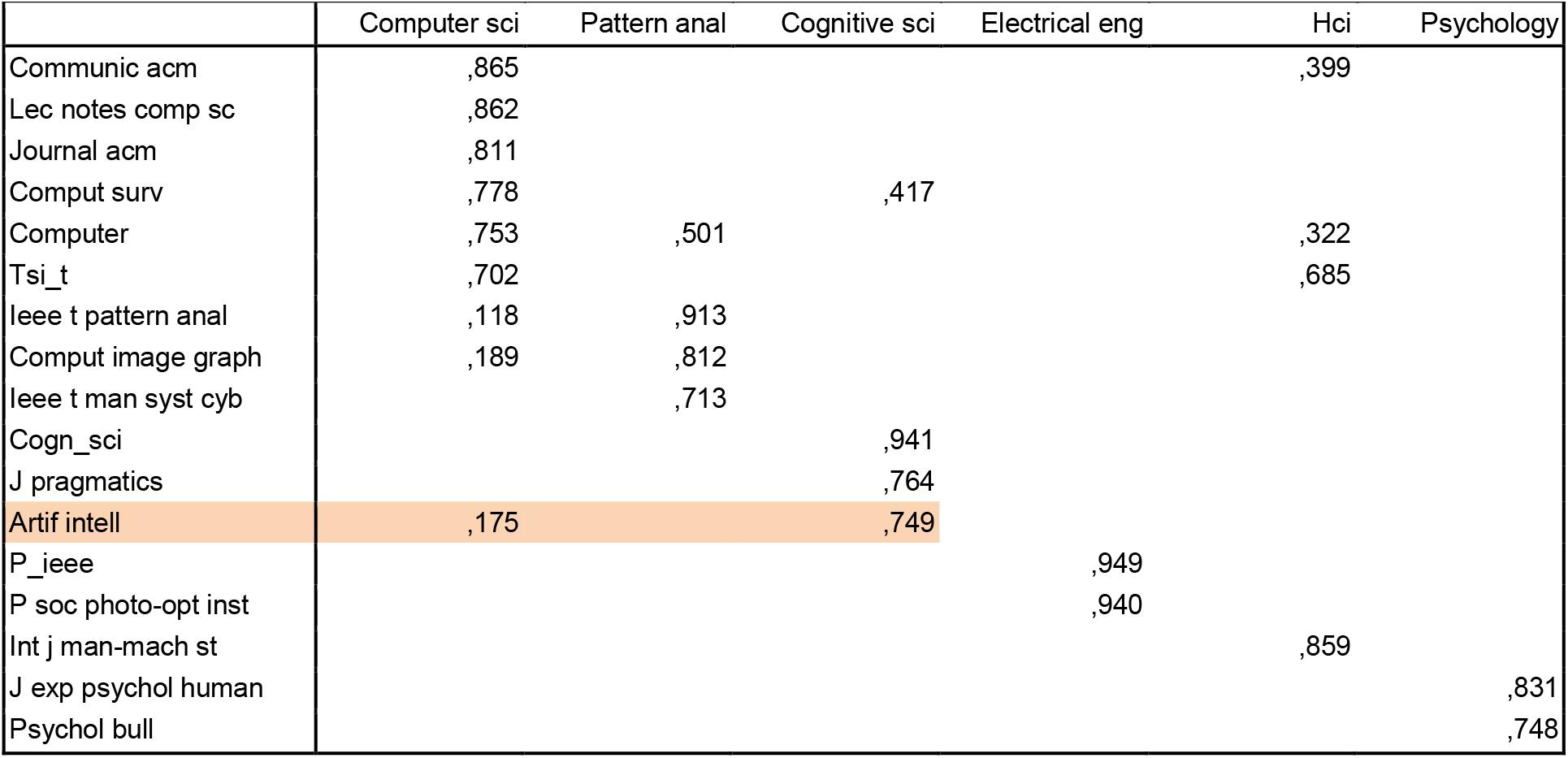
Artificial Intelligence (1982)

However, over the years the nature of AI radically changes. More and more, over the years the *Artificial Intelligence Journal* loads on a growing factor of mainly AI and applied AI journals. Increasingly this factor dominates its own environment, being an indicator for becoming more disciplinary (Van den Besselaar 2001): The AI factor becomes more homogeneous, and the journals generally have a *disciplinary citing pattern*, which means that the journals load on one factor only. Figure 6 shows this clearly for 2007.

AI is now the first factor in the citation network, and consists of AI journals mainly. Some multidisciplinary journals are still present in the AI factor, such as a robotics journal that also loads on the pattern recognition factor. More specifically, the environment of *Artificial Intelligence Journal* now is heavily computer science dominated, and much of the heterogeneous nature of its citation environment in the earlier period has disappeared.

With AI, we clearly have the second pattern in its dynamics: interdisciplinary fields emerge and over time develop into a mature state. There they have stabilized a citation network that similar to the ‘normal’ disciplinary pattern discussed above.

**Figure 6:**
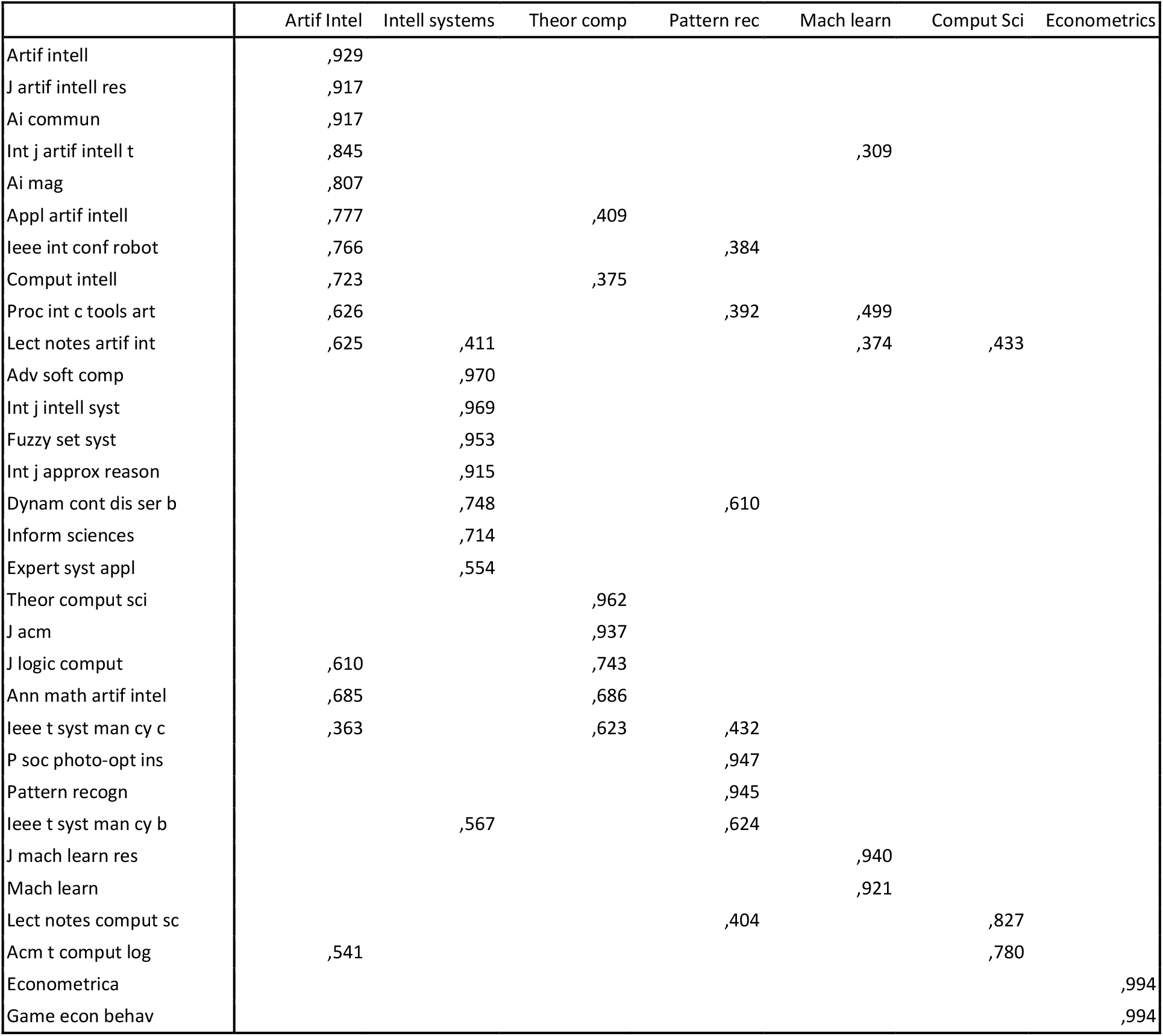
Artificial Intelligence (2007)

### 3.2 Cognitive science: Multidisciplinarity, failed interdisciplinarity and disappearing

Let’s move to another example: cognitive science. We analyze here the development of the journal *Cognitive Science* over the years, to inform us about the (multi)disciplinary nature of cognitive science research. Figures 7, 8 and 9 show the results of the factor analysis of the journal-journal citation matrix for the years 1988, 1998, and 2007. In the first year, we find a typical multidisciplinary pattern, with *Cognitive Science* loading lowest on two factors: cognitive psychology and artificial intelligence. This reflects the emphasis in cognitive science in those days on bridging between (and integrating) computer science (artificial intelligence) and psychology. In other words, the journal had a double identity. The larger environment of *Cognitive Science* consists of a variety of research fields within computer science and within psychology. How does cognitive science develop over the years? Ten years later, the pattern is more or less the same. The journal environment of *Cognitive Science* has become larger, but *Cognitive Science* is still in the tail of cognitive psychology and of computer science.

Again ten years later, in 2007, the picture has changed. *Cognitive Science* has a radically changed environment, as computer science related fields have become less central, and *Cognitive Science* has become part of the cognitive psychology factor. One may say that it has lost its computer science and artificial intelligence identity, and has been normalized within psychology. There is still knowledge exchange between computer science and cognitive science/psychology (not analyzed here), but the cognitive distance has become larger. In other words, the multidisciplinary interface between psychology and computer science, as embodied in journals like *Cognitive Science*, has disappeared.

These results provide us with the third model, emerging of multidisciplinary, unstable development and disappearance.

**Figure 7:**
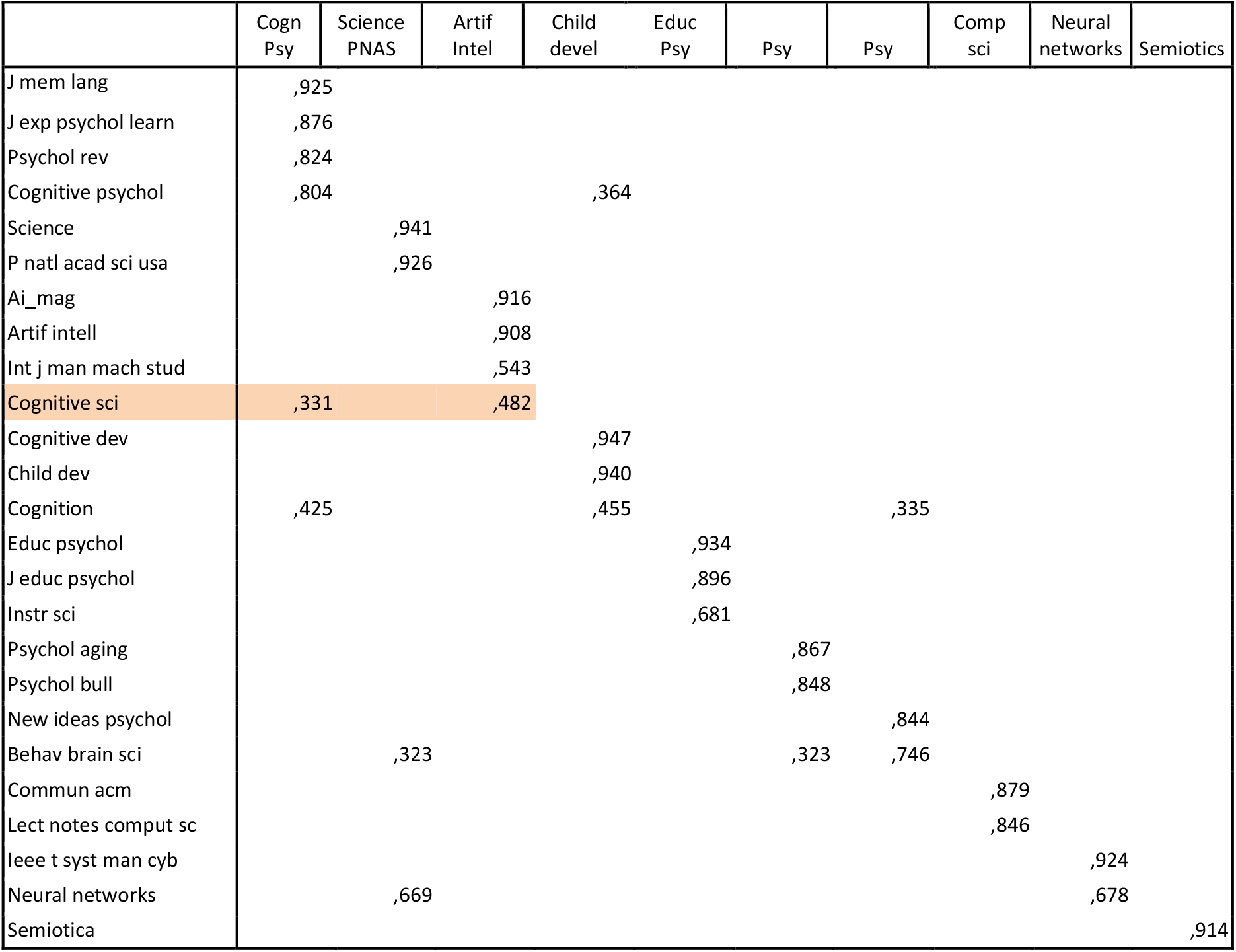
Cognitive science (1988)

**Figure 8:**
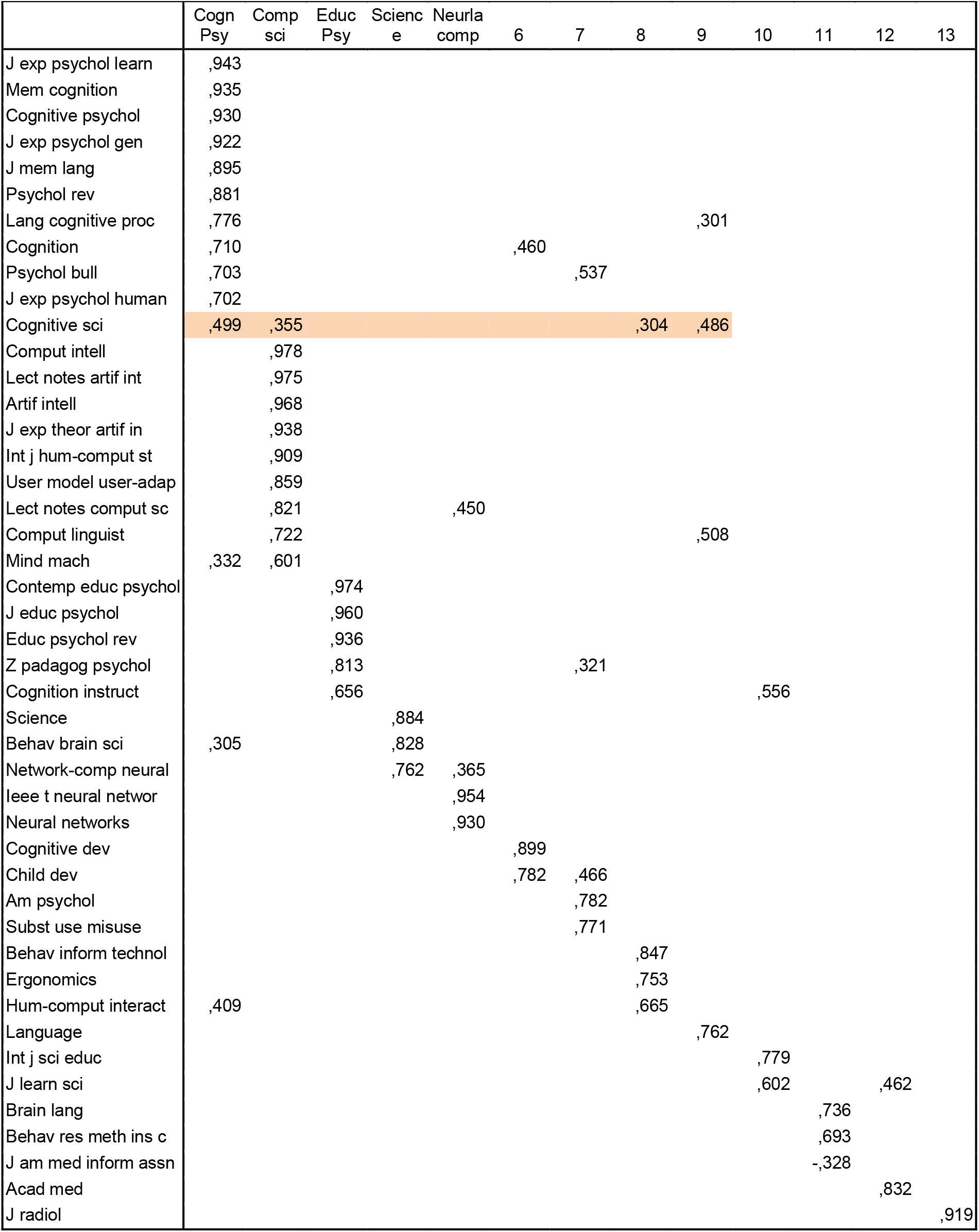
Cognitive science (1998)

**Figure 9:**
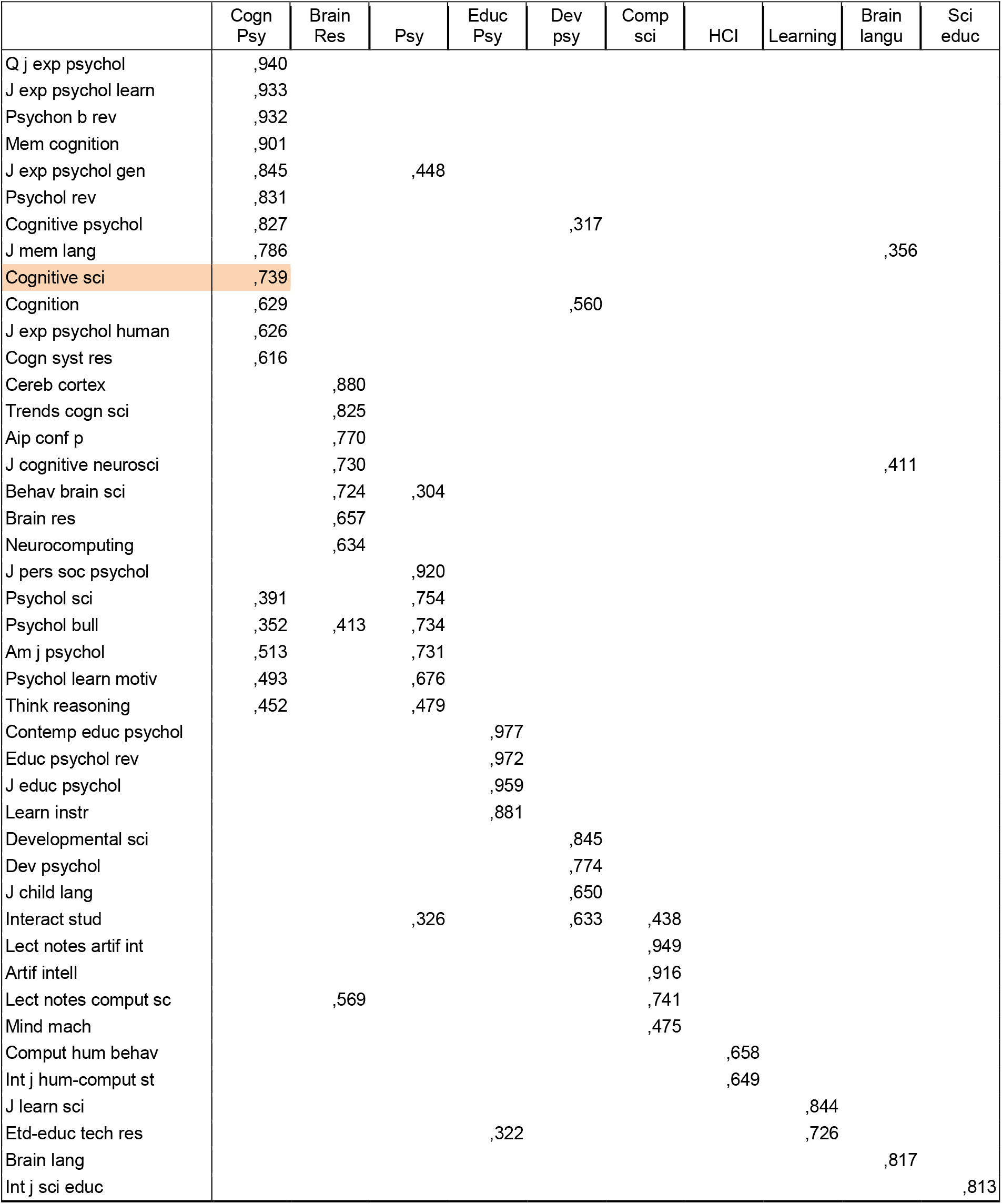
Cognitive Science (2007)

**Fig 9b:**
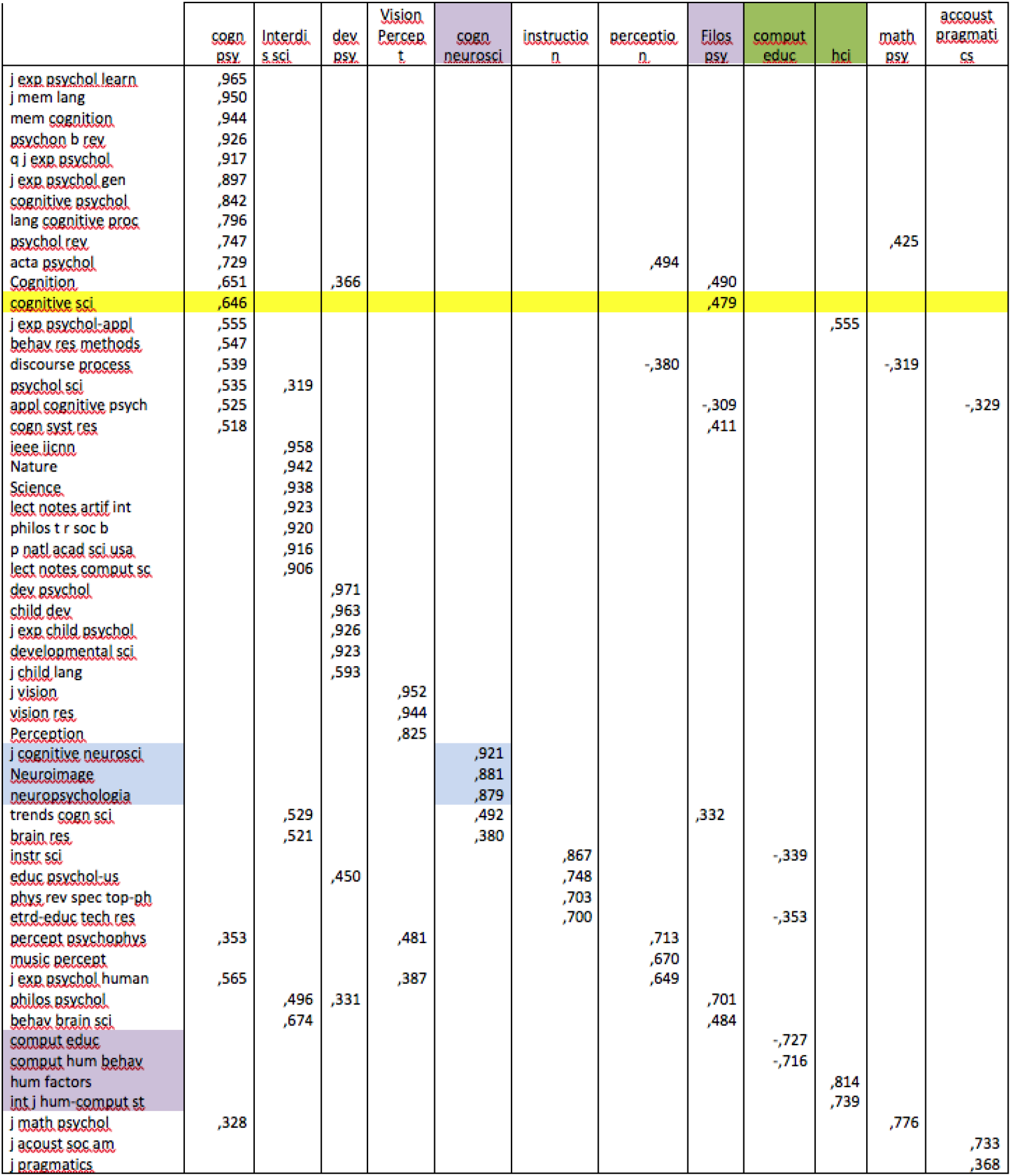
2008. Cognitive science also loads on a factor representing philosophy of prychology **Fig 3**: the citation network of cognitive science in 2008

However, a new research field has emerged in the environment of cognitive science / cognitive psychology, which is brain and neuroscience. As a test, we mapped the citation environment of a core journal of the newly emerged factor on brain and neuroscience: the *Journal of Cognitive Neuroscience*. The results are in figure 10. Interestingly, new multidisciplinary activities have developed, but now at the interface of cognitive neuropsychology and brain science, facilitated by new imaging technologies. The *Journal of Cognitive Neuroscience* is a main multi-identity journal that spans these boundaries.

## 4. The development of recent forms of interdisciplinarity

Does the approach also work for more recent fields? Here we try for genomics and for nanotechnology.

### 4.1 Genomics

We start with a list of the 23 main genomics journals, provided by Antoine Schoen (Paris-Est, IPTS). We determined the citation environment of the journals in 2007 and in 2000 (for those that already existed in 2000). In the following two figures we visualize the factor structure of genomics for the two years. The lines indicate the similarity between the journals (the nodes) in terms of citation behavior (so not: citation relations).

In 2000, the genomics journals were mainly in a factor with protein research and with bioinformatics. One of the genomics journals (*Genome*) loaded on the food research factor. Between 2000 and 2008 the genomics field did grow, but it also differentiated. In 2008 we therefore see quite a few factors with genomics journals. In other words, genomics is not a (inter)disciplinary research field, but it has differentiated into a series of fields applying genomics in existing research: (1) proteomics, (2) food genomics, (3) plant genomics, (4) human genomics, (5) genomics in cancer research, and (6) pharmarcogenomics.

**Figure 11:**
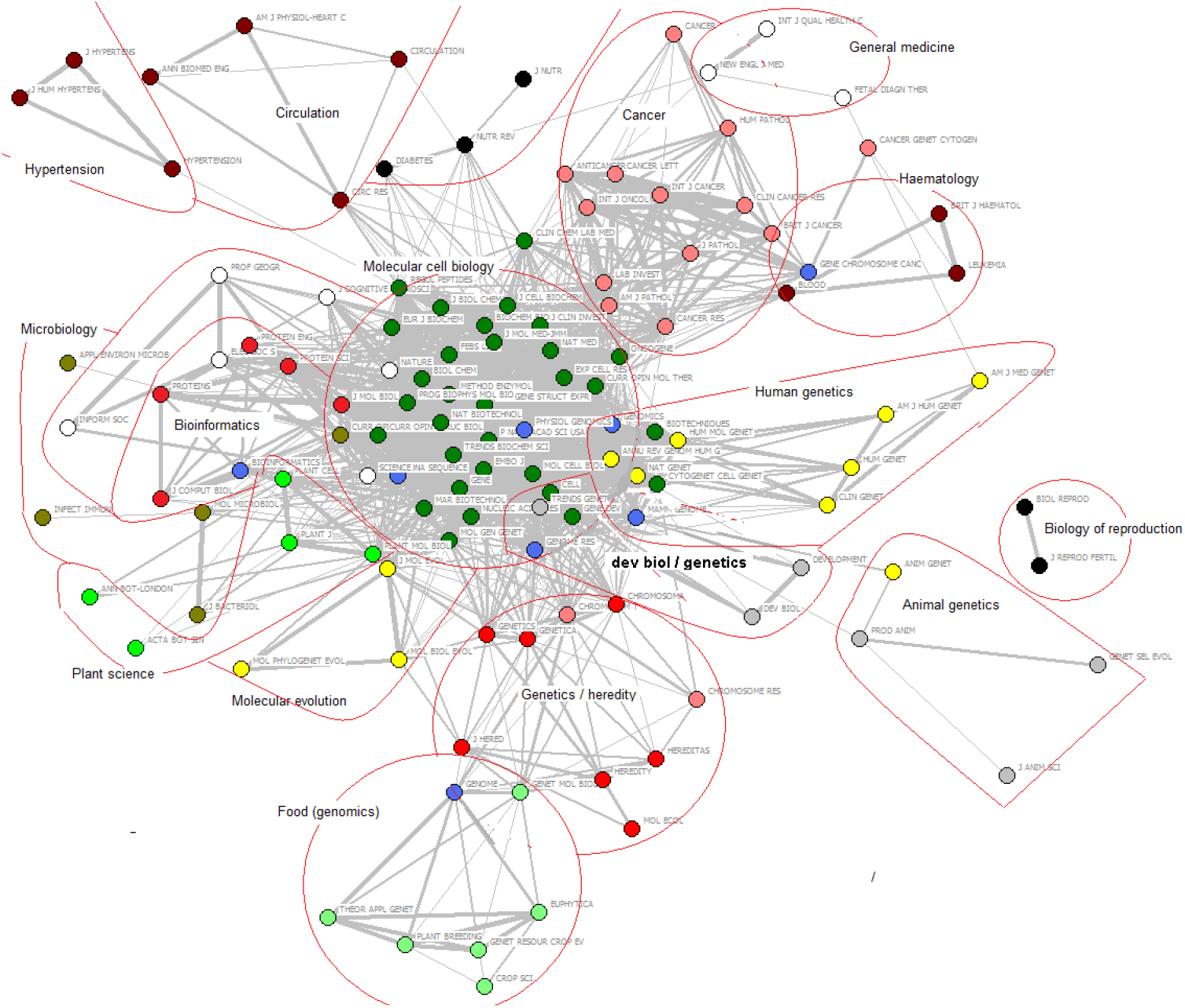
genomics 2000

**Fig 12:**
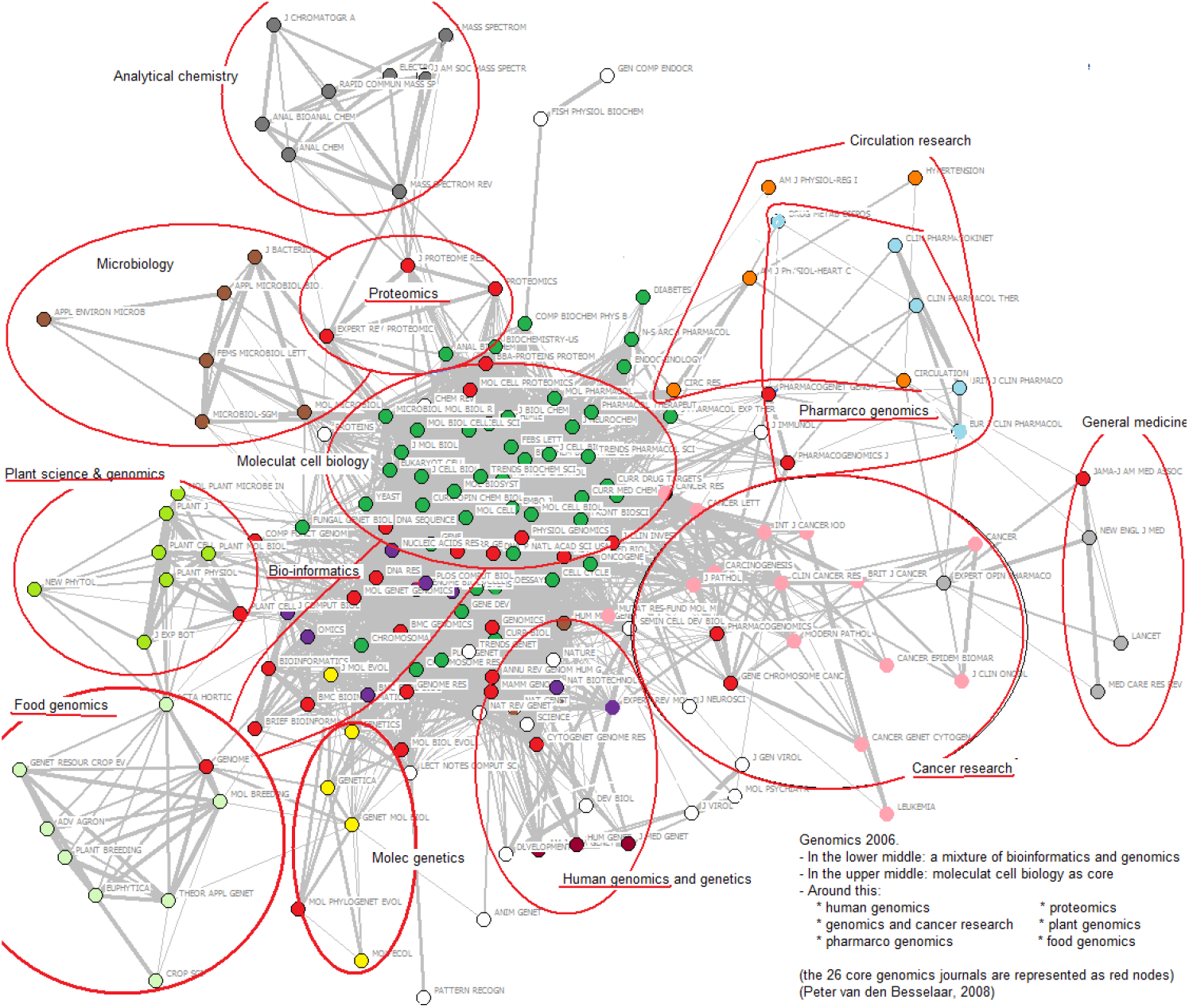
Genomics research (2007)

### 4.2 Nanoscience

The opposite development is visible in nanoscience, that emerged within chemical materials, applied physics, and in bionanoscience. However, using a journal-journal citation approach, a latent underling factor becomes visible over the years consisting of most nanoscience journals. This suggests that nanoscience merges in to a new research field with a ‘normal’ disciplinary communication (citation) pattern. At the same time, nanotech application in biomedicine and electronics are visible in other parts of the knowledge landscape.

## 5. Conclusions

We started the analysis with investigating the disciplinary journal citation network as a base line. As shown, disciplinary fields have a clear structure in of the journal-journal citation relations: the journals representing the disciplinary field under study all load on one single factor that represents the field.

In contrast to this disciplinary journal citation network, we distinguish a multidisciplinary one, which consists of journals with multiple identities: journals belonging to more research fields, and therefore loading on more than one factor – factors representing the fields the multidisciplinary journals relates to. As easy to understand (and as the factor loadings indicate), these multidisciplinary journals are not in the core of the fields, but more at the boundaries. Indeed, innovation within science happens at the boundaries between existing research fields and disciplines.

Using this finding, we could proceed in clarifying interdisciplinarity. Interdisciplinary fields, as we illustrated using of Artificial Intelligence as an example, are new developments emerging between fields, which over time stabilize and transform into ‘normal’ and ‘disciplinary’ research fields with similar citation structures/patterns as traditional disciplinary fields. In other words, there is no significant difference between disciplinary and *mature* interdisciplinary fields. Therefore, the focus on *change* in the journal citation networks is be more meaningful than discussions about *nature* interdisciplinary research – as it changes during its development. Change may mean a variety of things, such as <u>emerging</u> new fields, <u>growth</u>, <u>differentiation</u>, <u>merging</u>, <u>decline</u>, and <u>disappearance</u>. Elsewhere we will generalize this approach of interdisciplinary to knowledge dynamics in general (Van den Besselaar, Schoen et al – in preparation).

Of course, innovation may fail, and early multidisciplinary developments may not successfully develop into a new (first interdisciplinary and then new disciplinary) field. The example of cognitive science clearly indicates this. It continued to have an unstable position in the scholarly landscape, and more recently it was ‘absorbed’ by cognitive psychology.

Of course, one would like to understand under which conditions stabilization happens, and under which conditions it does not. And, are these conditions cognitive (the field is appealing enough to attract researchers; no competing fields emerge elsewhere), or social (funding for example, or the establishment of scholarly associations or of institutionalizing a field as a university program)? We cannot discuss this in depth here, but a few observations can start the process of understanding.

Firstly, the Artificial Intelligence case showed not only differentiation into a stable field, but also a relatively strong growth of the field. From year to year, more journals entered the field. Also, the field is in a stable way dominated by acknowledged core journals. In contrast, the cognitive science case is completely different. The field did not grow as interdisciplinary endeavor, but remained small between two growing related environments: (i) on the one hand the strongly growing field of disciplinary cognitive psychology, and (ii) the emerging AI field on the other. This may have reduced the cognitive space, hindering cognitive science to become a new interdisciplinary development. And more recently, a third competing field came up between cognitive psychology and biology: cognitive neuroscience came up. This all reduced the space for cognitive development, and the journal with that name remained therefore for several decades an isolated driving force. And obviously, without growth there is no possibility of stabilization of a new field.

If interdisciplinarity is a process – as we argued in this paper – the search and development of indicators for interdisciplinarity seems to be completely on the wrong track. More precisely, in the different phases of its development, interdisciplinarity has different characteristics. Indicators for should reflect this, and unfortunately, existing indicators do not do so.

Take for example the proposed interdisciplinarity indicator of *betweenness centrality* (Leydesdorff 2007)^10^. The idea is that the higher betweenness centrality, the more interdisciplinary a journal is. In the light of our results, this is a clear mistake as it actually only holds for early interdisciplinarity. The more the field grows and stabilized – that is, the more it is successful – the lower betweenness centrality will become. Consequently, high betweenness does not say anything about interdisciplinarity, if we do not know whether the field is in an early or late stage. The same holds for other indicators that recently have become popular recently, such as coherence and diversity (Rafols & Meyer 2010). Also these indicators have values related to the phase of development the interdisciplinary field is in. Interdisciplinary success here means actually increasing disciplinarity.

Quantitative indicators for interdisciplinarity should therefore measure at least three things: (i) the age and phase of a new field, partly indicated by growth or lack of it; (ii) the level of multidisciplinarity the of the involved journals, that is the structure of the citation network of the field under study, and (iii) the relations with other fields in the environment. With a set of indicators like this, we are able to identify interdisciplinary forms of knowledge dynamics, but also other forms.

Summarizing, interdisciplinarity should be considered as a specific form knowledge dynamics, and not as a characteristic of a journal or field.

**Table 10.** *Journal of cognitive neuroscience (2006)*- 8 factor solution

|  | Neuro-Science | Cogn<br>Neuro<br>Psychol | Cognitive<br>psychology | Science/<br>Nature | Brain<br>research | Human<br>brain<br>mapping | Psycho-<br>physiol | Percep-<br>tion |
| --- | --- | --- | --- | --- | --- | --- | --- | --- |
| J neurosci | .977 |  |  |  |  |  |  |  |
| Curr opin neurobiol | .959 |  |  |  |  |  |  |  |
| Trends neurosci | .959 |  |  |  |  |  |  |  |
| Annu rev neurosci | .946 |  |  |  |  |  |  |  |
| Nat neurosci | .939 |  |  |  |  |  |  |  |
| Eur j neurosci | .916 |  |  |  |  |  |  |  |
| Neuron | .906 |  |  |  |  |  |  |  |
| Nat rev neurosci | .900 |  |  | .366 |  |  |  |  |
| Cereb cortex | .860 |  |  |  |  | .357 |  |  |
| Neuroscience | .820 |  |  |  | .455 |  |  |  |
| J physiol-paris | .791 |  |  | .349 |  |  |  |  |
| J neurophysiol | .761 |  |  |  |  |  |  |  |
| Brain res | .712 |  |  |  | .540 |  |  |  |
| Neuroreport | .698 |  |  |  | .332 | .437 |  |  |
| Neurosci biobehav r | .682 |  |  |  | .483 |  |  |  |
| Neurosci lett | .671 |  |  |  | .578 |  |  |  |
| Trends cogn sci | .627 |  |  | .336 |  | .361 |  |  |
| Neuropsychology |  | .906 |  |  |  |  |  |  |
| J int neuropsych soc |  | .902 |  |  |  |  |  |  |
| Cortex |  | .856 |  |  |  |  |  |  |
| Brain cognition |  | .825 |  |  |  | .352 |  |  |
| Neuropsychologia |  | .824 |  |  |  | .375 |  |  |
| Cogn neuropsychol |  | .675 |  |  |  |  |  |  |
| Brain lang |  | .598 |  |  |  |  |  |  |
| Brain |  | .593 |  |  | .436 |  |  |  |
| Neurology |  | .445 |  |  | .419 |  |  |  |
| J exp psychol gen |  |  | .923 |  |  |  |  |  |
| J exp psychol learn |  |  | .922 |  |  |  |  |  |
| Mem cognition |  |  | .897 |  |  |  |  |  |
| J mem lang |  |  | .874 |  |  |  |  |  |
| Psychol rev |  |  | .786 |  |  |  |  |  |
| Psychol sci |  |  | .681 |  |  |  |  | .334 |
| Cognition |  |  | .558 | .329 |  |  |  |  |
| Lect notes comput sc | .313 |  |  | .896 |  |  |  |  |
| Science |  |  |  | .862 |  |  |  |  |
| P natl acad sci usa | .379 |  |  | .862 |  |  |  |  |
| Nature | .300 |  |  | .835 |  |  |  |  |
| Ann ny acad sci | .359 |  |  | .806 |  |  |  |  |
| Prog brain res | .506 |  |  |  | .692 |  |  |  |
| Brain res bull | .598 |  |  |  | .690 |  |  |  |
| Exp brain res | .336 |  |  |  | .500 |  |  | .478 |
| Clin neurophysiol |  |  |  |  | .466 |  | .428 |  |
| Hum brain mapp |  |  |  |  |  | .897 |  |  |
| Neuroimage |  |  |  |  |  | .880 |  |  |
| J cognitive neurosci | .332 | .513 |  |  |  | .672 |  |  |
| Cogn affect behav ne | .511 | .380 |  |  |  | .569 |  |  |
| Psychophysiology |  |  |  |  |  |  | .956 |  |
| Boil psycho |  |  |  |  |  |  | .949 |  |
| Int j psychophysiol |  |  |  |  |  |  | .917 |  |
| Percept psychophys |  |  | .372 |  |  |  |  | .766 |
| Vision res |  |  |  |  |  |  |  | .687 |
| J exp psychol human |  |  | .603 |  |  |  |  | .664 |

## Footnotes

1 This paper was written in 2012, and is now being updated with more recent data on the developments of the fields we use as cases here: cognitive science, genomics and a few others. It is based on an approach of interdisciplinarity not as a characteristic of research fields or disciplines, but as a characteristic of the dynamics of disciplines. A first formulation is in Van den Besselaar & Heimeriks 2001, and it was applied in several case studies, among those Vugteveen et al 2014.

2 A special case of this interest, of course, is ‘converging technologies’ (nano, bio, info, cogno) which is supposed to be the revolution of the future.

3 Often it is argued that with online open access, scholars do not use journals any more for finding the material they consider relevant, but directly access articles. If this is the correct, one would expect different patterns of journal-journal citations nowadays than earlier. This does not seem to be the case, suggesting the role journals remain playing.

4 If journals exist, we use these for mapping the development of a field, and use papers within the determined journal set to map research fronts within the field. In case of newly emerging fields, where no dedicated journals exist yet, we use a key word based approach and identity in a second step the most prominent journals in the retrieved corpus of papers. These can be used for an initial analysis at the journal level. See: Peter van den Besselaar & Thomas Gurney 2008.

5 These are often not clearly distinguished, resulting into wrong conclusions, e.g., on cognitive science (Goldstone & Leydesdorff 2006)

6 For details about the method: Van den Besselaar, Leydesdorff 1996; Van den Besselaar, Heimeriks 2001.

7 The main journal is here defined as the (non-review) journal with the most citations in the field. We do not go for the impact factor, as size matters for impact.

8 The following analysis is not dependent on the choice of this journal. For chemistry as a whole, using the main journals in a variety of chemical fields for determining the journal citation network of chemistry: organic chemistry, inorganic chemistry, biochemistry, physical chemistry, chemical materials, catalysis. Although the resulting network is much bigger, this does not influence the argument made here. We also redid the analysis using one or more entrance journals of each of the fields within chemistry. This creates maps of the specific fields, which do not differ from the broad picture emerging here.

9 Using a threshold of 0.5%. This means we include all journals that are responsible for at least 0.5% of the citations to the *JACS*, and all journals that are good for at least 0.5% of the references in the *JACS*.

10 The ‘betweenness centrality’ indicator is also confusing two other reasons. (i) Betweenness centrality is a *relational* concept that refers to centrality in bridging communication. But it is used for a *positional*phenomenon, identifying similar positions in the citation network. A good example where it goes wrong is an analysis of cognitive science (Goldstone and Leydesdorff 2006), where the authors confuse the relational role of the journal *Cognitive Science* with its position in the citation network. (ii) The indicator is used as if the citation network is unvalued, as it translates a valued network (journals as nodes, correlations as links) into an unvalued one, by introducing a threshold.

